# Natural variation in growth and physiology under salt stress in rice: QTL mapping in a *Horkuch* × *IR29* mapping population at seedling and reproductive stages

**DOI:** 10.1101/2020.03.01.971895

**Authors:** Taslima Haque, Sabrina M Elias, Samsad Razzaque, Sudip Biswas, Sumaiya Farah Khan, G.M. Nurnabi Azad Jewel, Md. Sazzadur Rahman, Thomas E. Juenger, Zeba I Seraj

**Affiliations:** Plant Biotechnology Lab, Department of Biochemistry and Molecular Biology, University of Dhaka, Dhaka-1000, Bangladesh; Department of Integrative Biology and Institute for Cellular and Molecular Biology, University of Texas, Austin, Texas 78712, USA; Plant Physiology Division, Bangladesh Rice Research Institute, Gazipur, Bangladesh; Department of Agronomy and Horticulture, University of Nebraska, Lincoln, Nebraska 68583, USA; Department of Biochemistry and Molecular Biology, Jagannath University, Dhaka-1100, Bangladesh; School of Life Science, Independent University, Dhaka-1229, Bangladesh

**Keywords:** salinity, QTL, reciprocal cross, cytoplasm, crop breeding

## Abstract

Salinity has a significant negative impact on production of rice. To cope with the increased soil salinity due to climate change, we need to develop salt tolerant rice varieties that can maintain their high yield. Rice landraces indigenous to coastal Bangladesh can be a great resource to study the genetic basis of salt adaptation. In this study, we implemented a QTL analysis framework on a reciprocal mapping population between a salt tolerant landrace *Horkuch* and a high yielding rice variety *IR29*. Our aim was to detect genetic loci that contributes to the salt adaptive responses of the two different developmental stages of rice which are very sensitive to salinity stress. We identified 14 QTL for 9 traits and found that most are unique to the specific developmental stage. In addition, we detected a significant effect of the cytoplasmic genome on the QTL model for some traits such as leaf total potassium and filled grain weight. This underscores the importance of considering cytoplasm-nuclear interaction for breeding programs. Along with this, we identified QTL co-localization for multiple traits that highlights the possible constraint of multiple QTL selection for breeding programs due to different contributions of a donor allele for different traits.

**Highlights:** We identified genetic loci for the salt tolerance response of two different developmental stages of the rice plant and detected significant contribution of cytoplasm-nuclear genome interaction for a few traits.

## Introduction

Rice (*Oryza sativa* L.) production, which feeds almost half of the world population, is under threat from global environmental changes such as increasing salinity, heat and drought (Seck et al., 2012; Ashikari and Ma, 2015). Among these abiotic stresses, salinity has already affected 45 million hectares of irrigated land worldwide and 1.5 million additional hectares are impacted each year (Munns and Tester, 2008). Bangladesh and other locales at or near sea level are particularly vulnerable to climate change-induced salinity. In Bangladesh, about 30% of the cultivable land along the coast is affected by salinity due to tidal flood during the wet season resulting in direct inundation by saline water, and upward or lateral movement of saline ground water during the dry season (Haque, 2006).

High yielding “elite” rice cultivars are especially susceptible to salinity stress. Recent studies have shown that production of high yielding rice varieties in Bangladesh will decrease by 15.6% in coastal districts where soil salinity is predicted to exceed 4 deciSiemens per meter (dSm^−1^) by 2050 (Dasgupta *et al*., 2014). However, the coastal belt of Bangladesh is enriched with many local rice landraces, among which a handful are adapted to high-to-moderate soil salinities. The rice landrace, *Pokkali*, has long been used as a salt tolerant landrace reference. Many other salt-tolerant landraces such as *Horkuch, Ashfal, Jatai* and *Balam* from the coastal region of southern Bangladesh have been identified and are currently grown by farmers in these salt-affected regions (Lisa *et al*., 2004; Rahman *et al*., 2016). Unfortunately, these landraces suffer from low yield, poor grain quality and longer duration to reach maturity and therefore cannot serve as good candidates for commercial crop varieties. However, studies of these adapted landraces to understand their salt tolerant mechanisms can open opportunities to incorporate desired traits to commercial rice varieties. Therefore, it is important for breeders to identify genetic variants of salt stress responses in these adapted landraces in order to design highly salt tolerant rice.

The effect of salinity on rice growth varies across various developmental stages (Lutts *et al*., 1995). The rice plant is most sensitive to salinity at the early seedling stage and during panicle formation, whereas it is relatively tolerant during early germination, active tillering and maturity (Akbar *et al*., 1972; Heenan *et al*., 1988; Lutts *et al*., 1995; Pearson and Bernstein, 1959; Singh and Flowers, 2010; Zeng and Shannon, 2000). During salt stress at the early seedling stage, there is a significant decrease of dry matter as well as quantum yield of PSII and a significant increase of sodium concentration in root, stem and shoot tissue (García Morales *et al*., 2012). At the reproductive stage, physiological studies under salinity stress show a significant decrease in panicle weight, panicle length, primary branches per panicle, filled grains per panicle, total seeds per panicle, total seed weight per panicle, 1000-seed weight and total seed weight per plant (Abdullah *et al*., 2001; Rao *et al*., 2008). Moradi et al. (2003) have however shown that salinity tolerance at the seedling and reproductive stages is only weakly associated. This emphasizes the importance of discovering the contributing traits of these two very important growth stages of rice.

The physiological basis of salt tolerance during the early seedling stage is well understood. Munns and Tester (2008) and Roy (2014) have proposed several physiological mechanisms of seedling tolerance such as sodium exclusion, compartmentalization of excessive sodium ions (tissue tolerance) and shoot-ion independent tolerance for early stage tolerance. It has been reported that *Pokkali* maintain lower shoot Na^+^ accumulation and lower shoot Na^+^/K^+^ ratio under high salinity compared to sensitive genotypes (Kavitha *et al*., 2012; Sexcion *et al*., 2009). The enhancement of salinity tolerance by constitutive overexpression of the vacuolar Na^+^/H^+^ antiporter gene from *Pokkali* in transgenic rice plants suggest that this landrace may use a tissue tolerance mechanism to lower shoot Na^+^/K^+^ ratio under high salinity (Amin *et al*., 2016). Negrao et al genotyped 392 rice accessions by EcoTILLING in order to understand allelic difference for salt stress. They targeted five known genes that are involved in these different salt tolerant mechanisms and assembled a set of accessions that represents all the haplotypes present in the coding region of these five genes. The systematic study of phenotypes of this set suggest that none of the main three mechanisms of tolerance is preferentially used over another (Inês *et al*., 2015). Therefore, studies of different landraces can offer ways to understand individual salt tolerance mechanisms in the rice plant (Lisa *et al*., 2004; Yesmin *et al*., 2014). However, the mechanism associated with tolerance during the reproductive stage has been barely explored. As mentioned earlier, high salinity in this stage can alter many traits associated with grain quality and quantity, eventually decreasing yield significantly. Therefore, it is important to explore the physiological response of the rice plant at both stages in order to obtain a superior variety from breeding which can maintain salinity tolerance for both developmental stages.

The choice of female parents in breeding programs plays a critical role in the performance of crosses. Plants show evidence for complex nuclear-cytoplasmic interaction that may alter their phenotypes in both interspecific and intraspecific crosses. However, it still remains unclear to what extent these two components interact with each other and the role of environment in this interaction. Gregorio and Senadhira (1993) have studied the genetics of salinity tolerance on diallelic reciprocal crosses of nine different rice varieties and found significant reciprocal effects among crosses. The presence of maternal inheritance has also been reported for other abiotic stresses such as chilling response (Chung et al., 2003) and drought (Iida *et al*., 2007). Therefore, in plant breeding programs where the aim is to produce stress tolerant high-yielding varieties, it is important to consider a specific cytoplasm and its interactions with nuclear donor alleles for determining the performance of plants under stress.

Quantitative trait loci (QTL) mapping has been implemented in many studies of rice to explore the genetic basis of traits involved in salinity stress for seedling stages, including salt injury/tolerance score, fresh and dry weight of shoot and root, Na^+^ and K^+^ content of shoot and root, and chlorophyll content (Cheng *et al*., 2011; Lin *et al*., 2004; Ren *et al*., 2005; Sabouri *et al*., 2009; Soltani *et al*., 2016; Thomson *et al*., 2010; Tian *et al*., 2011; Wang *et al*., 2012; Zheng *et al*., 2015). However, very few studies have been conducted to understand the genetic basic of reproductive stage traits that are important for tolerance such as plant height, tiller number, panicle number, pollen fertility and yield (Hossain *et al*., 2015; Zang *et al*., 2008).

QTL co-localization has been reported for traits that are strongly correlated (Dechaine *et al*., 2014). Many clustered, putatively pleiotropic QTL have been found that affect various life history and fitness characters, especially those that are related to yield, in rice, wheat, pea and rapeseed (Burstin *et al*., 2007; Quarrie *et al*., 2006; Shi *et al*., 2009; Xue *et al*., 2008). QTLs for two different traits can have the same/opposite sign of effects for both. If the genes involved help in coordination during multiple steps of development, then positive selection for one trait may have an outcome on several traits in the same positive direction, e.g. a pleiotropic same sign QTL for seed size and protein content in pea (Burstin *et al*., 2007). Very strong opposite sign of effects for QTL has also been reported for plants including rice (Dechaine *et al*., 2014; Xiao *et al*., 1998) and may represent tradeoffs. For breeding programs where breeders aim for QTL pyramiding for multiple desired traits, opposite signed QTL for different traits may impose some constraints on selecting co-localized QTL. Hence, it is beneficial to have a clear understanding about QTL co-localization and perform careful selection of these genomic loci for pyramiding.

In this study we genotyped a reciprocal mapping population between the salt tolerant landrace, *Horkuch* and a high yielding variety *IR29* by DArTseq technique (Akbari *et al*., 2006). We identified 14 QTL for 9 traits for salinity treatments at two different developmental stages of the rice plant. One important finding of this study was to characterize the role of cytoplasm in a plant’s performance under salinity and implement an analysis incorporating this information to estimate the effect of a QTL. Furthermore, in this QTL analysis framework, we applied a linear mixed model to incorporate residual polygenic variation which is a better way to estimate QTL effect for polygenic traits. In our previous study we applied Double digested Restriction Associated DNA (ddRAD) technique to construct genetic map of this population where we failed to map a substantial genetic space (Noor *et al*., 2019). In this current study, with the aid of an improved genetic map by DArTseq technique and a robust QTL analysis framework we were able to identify additional QTLs with higher likelihood and tighter confidence interval. In addition to that, we identified co-localized QTL within and across two different treatment stages, which emphasizes the need for conditional selection of QTL in a breeding program in order to combine survivability at the seedling stage and yield tolerance at reproductive stage. Taken together, the findings of this study contribute to our understanding of the molecular mechanism of salt tolerance for *Horkuch* and pave a way to introgress salinity tolerance into a commercial cultivar that can maintain significant yields under stress.

## Methods

### Development of the reciprocally crossed populations and physiological screening

The rice cultivars *Horkuch* (IRGC 31804) and *IR29* (IRGC 30412) were used as parents to raise a bi-parental reciprocal mapping population. Detailed method for developing this mapping population and physiological screening for different developmental stages can be found in Elias et al (2020). In brief, our experimental approach centers on an F_2:3_ design, whereby genotypes for mapping are collected from F_2_ individuals and phenotypes are obtained from a sampling of their F_2_:_3_ progenies (Zhang et al, 2004). In this article, F_2:3_ progenies derived from *Horkuch* (mother) × *IR29* (father) will be referred to as *Horkuch*♀ and those from *IR29* (mother) x *Horkuch* (father) as *IR29*♀. We randomly chose 137 families from *IR29*♀ and 65 families from *Horkuch*♀ cross for seedling stage QTL analysis. For reproductive stage QTL analysis 140 families were chosen based on a selection of 70 F_2_ families from each population. Our selection was based on the distribution of SES scores where the lower tail (more tolerant families) was defined as SES scores from 3 to 5 and the upper tail (sensitive) was defined as SES scores from 7 to 9. All families that were in the lower or upper tail were selected along with 70 randomly chosen families from SES score in between 5 and 7. The two F_0_ parents were also included in our studies. A few families had poor germination and were subsequently excluded from the reproductive screening experiment. In the end, 130 families were included in reproductive screening: 61 from *Horkuch*♀ and 69 from *IR29*♀ population.

### DNA extraction, genotyping by DArTseq and linkage map construction

Genomic DNA of F_2_ individuals and parents was extracted using the CTAB method (Doyle and Doyle, 1990) from 1g of fresh leaf tissue after freezing in liquid nitrogen and grinding. Genotyping was done by the DArTseq technique as described by Akbari et al (2006). For the DArTseq method Nipponbare Genome from Phytozome (v9) was used as reference to determine the physical position of each DArTseq clone. We have filtered all the DArTseq SNPs and retained only loci that: a) are homozygous for both the parents and b) are polymorphic between parents. Analyses for linkage map construction were completed with qtlTools (Lovell, 2016) and R/qtl (Broman *et al*., 2003) packages. We filtered markers that had more than 50% missing data and showed significant segregation distortion for chi-square test (p-value < 0.001) from expected ratio (For a given locus,1:2:1 ≡ homozygous parent 1 allele: heterozygous: homozygous for parent 2 allele). Similar markers were further removed using dropSimilarMarkers function using qtlTools package (Lovell, 2016) in R with a minimum recombination fraction threshold of 0.03. Marker order was obtained by the tspOrder function from TSPmap tool (Monroe *et al*., 2017) which applies a traveling salesperson problem solver to order markers using Hamiltonian circuit. Markers which had discordance (have very different orders in genetic map vs. physical map) were also removed. Before estimating the linkage map a few alleles that showed erroneous call were masked manually. The final linkage map was estimated by est.map function of R/qtl package with the Kosambi map function using error probability threshold of 0.001. To test for the occurrence of cytoplasm-nuclear association, we performed chi-square test of independence on allele frequency for each locus grouped by cytoplasm and significant association was determined by adjusting p-values using FDR method (FDR threshold =0.1).

### QTL analysis

Genotype probabilities were calculated at a 1 cM step interval using the calc.genoprob function. QTL mapping was executed using the Haley-Knott regression algorithm (Broman *et al*., 2003) of the R/qtl2 (Broman *et al*., 2019) package where we fit the LOCO (Leave One Chromosome Out) model for each trait and included kinship as a covariate. The LOCO model utilizes the kinship matrix to reduce background polygenetic variation except for the chromosome that is being tested for QTL mapping. We also tested cytoplasm as an additive and interactive covariate in the QTL model using likelihood ratio tests and retained factors in the models when significant. To test whether our selection of cohorts imposed significant population structure for the reproductive stage treatment we assigned each F_2_ family into one of the following categories: tolerant, intermediate and sensitive. We incorporated selection cohort as a covariate while building the primary QTL model for traits at reproductive stage. However, we did not find any significant effect of selection cohort on QTL models and therefore did not include selection cohort as a covariate for further analysis. Significance thresholds for QTL were determined for each trait by 1000 permutations (alpha=0.05) and QTL peaks that passed the threshold were considered for further analysis. Permutations were stratified by cytoplasm for the QTL models where cytoplasm was considered as a covariate. We also evaluated the normality of the QTL model residuals and found this assumption was violated for the trait TK. Confidence intervals (1.5 LOD drop) for each QTL were calculated using the *lodind* function of the R/qtl package (Broman *et al*., 2003) expanding it to a true marker on both sides of the QTL. Codes for QTL analysis are available in GitHub repository (Haque, 2019).

### Identification of candidate genes within QTL confidence interval and Gene ontology enrichment analysis

In order to identify candidate gene models within a given QTL interval, we integrated the genetic and physical maps based on the marker order of the genetic map. We first pulled out the genetic markers flanking a given QTL confidence interval and their basepair positions to define the physical interval on the genome for that QTL. Gene models in these physical QTL intervals were retrieved using the structural gene annotation of the rice Nipponbare reference genome from Phytozome 9. We used the Gene ontology (GO) annotated for each gene model of this reference (Phytozome 9) for GO enrichment analysis. We then tested for the enrichment of GO terms for each QTL interval using the classical Fisher’s exact test available in the topGO (Alexa and Rahnenfuhrer, 2019) package in R.

## DATA AVILABILITY

Data for genetic map and phenotypes is available in GitHub repository (Haque, 2019)

## RESULTS

### Phenotypic traits vary between cross direction in both developmental stages

In our previous study, we used this bi-directional F_2_:_3_ experimental design on to examine the effect of salinity on various growth, yield and physiological parameters of rice as well as the role of cytoplasm on these traits and reported that maternal inheritance contributed to salt tolerance for F_3_ progenies (Elias *et al*., 2020). Rice is most susceptible to salinity during seedling and reproductive growth stages. Therefore, in the current study, we focused on such reported phenotypes from Elias et al (2020) that can potentially mediate stress during salinity treatment at these two developmental stages and the effect of cytoplasm-nuclear interaction. For the seedling stage, we evaluated traits that were related to survival, photosynthesis and mineral elements in leaves including Standard Evaluation Score (SES), Total Chlorophyll Content (Tchlr), Shoot Length (SL), Root length (RL), Total Sodium (TNa), Total Potassium (TK) and Potassium by Sodium (K/Na). For the reproductive stage treatment, we focused on yield-related parameters including Panicle Exsertion (PE), Total Tiller Number (TT), Effective Tiller Number (ET), Filled Grain Weight (FGW), Filled Grain Number (FGN) and Spikelet Fertility (SF).

We found striking difference between the two parents for many of our measured traits across both stages of salt treatment (Supplementary Table 1). As reported in previous studies, we have found that *Horkuch* is more tolerant to salinity (measured by SES score) compared to *IR29* which is highly sensitive for salt at the seedling stage (p-value < 0.01). For reproductive stage salinity treatment, these two parents show significantly different responses for ET, SF, FGW, DF and HI where the *Horkuch* parent had higher number of ET and also higher SF, FGW and HI. It has been reported that complex cytoplasm-nuclear interaction can alter plant phenotypes (Joseph *et al*., 2013a). We found many traits were significantly different as a result of cytoplasmic background (Supplementary Table 1). Tchlr, TNa, TK and K/Na were found to differ significantly among the *Horkuch*♀ and *IR29*♀ at the seedling stage treatment. For reproductive stage treatment PE, TT, ET, FGN were found to differ significantly among the two parental cytoplasm. This observation indicates that cytoplasm can explain a significant amount of variation in the population.

**Table 1:** QTL models

| Phenotypes | QTL model | Chr | Position | LOD | Lower CI | Upper CI | Positive Allele |
| --- | --- | --- | --- | --- | --- | --- | --- |
| SL | qSL.1@183 | 1 | 183.00 | 13.42 | 175.11 | 190.57 | Horkuch |
| SL | qSL.3@218 | 3 | 218.00 | 4.82 | 212.12 | 236.45 | Horkuch |
| SL | qSL.5@160 | 5 | 160.43 | 3.83 | 102.86 | 170.27 | Horkuch |
| RL | qRL.2@167 | 2 | 167.00 | 10.67 | 161.14 | 176.67 | Horkuch |
| TK | qTK.2@45* Cyto | 2 | 45.00 | 6.11 | 24.05 | 66.99 | Horkuch<sup>†</sup> |
| TK | qTK.3@204* Cyto | 3 | 203.78 | 6.46 | 194.44 | 209.29 | Horkuch<sup>†</sup> |
| PH | qPH.1@215 | 1 | 215.00 | 5.59 | 175.11 | 222.54 | Horkuch |
| PH | qPH.3@211 | 3 | 211.02 | 5.15 | 203.78 | 272.38 | Horkuch |
| PH | qPH.5@144 * Cyto | 5 | 144.00 | 6.64 | 124.73 | 170.27 | Horkuch<sup>†</sup> |
| ET | qET.7@97 * Cyto | 7 | 97.00 | 5.82 | 85.83 | 104.41 | Horkuch |
| FGN | qFGN.10@58 * Cyto | 10 | 58.48 | 7.72 | 50.30 | 107.07 | IR29 |
| FGW | qFGW.10@58 * Cyto | 10 | 58.48 | 9.13 | 50.30 | 107.07 | IR29 |
| SF | qSF.10@59 * Cyto | 10 | 59.00 | 7.71 | 50.30 | 107.07 | IR29 |
| HI | qHI.10@104+ Cyto | 10 | 103.75 | 8.48 | 50.30 | 107.07 | IR29 |

Each QTL model was built by linear mixed model using kinship matrix as a covariate. \*cytoplasm denotes interaction whereas (+) sign denotes only additive cytoplasmic effect in the QTL model. † denotes QTL that has both main and interaction effect since only considering direction of the main effect can be misleading

The generation of extreme phenotypes in a crossing population, or transgressive segregation, has been reported in many plants which may be due to the effect of complementary genes, over-dominance or epistasis (Rieseberg *et al*., 1999). Here, we further investigated the role of cytoplasm on the transgressive segregation of these traits. In this experiment we found that SES, SRWC, TChlr, TK, FGN, FGW clearly segregate transgressively (Supplementary Table 1). Interestingly, TK, TChlr, FGN and FGW showed significant differences in the two reciprocal crosses with respect to the segregation pattern. The distribution of TK showed a strong bimodal pattern with very little overlap of distribution between cytoplasmic backgrounds (Figure 1). This observation is indicative of an important role of cytoplasm on transgressive segregation for this population.

**Figure 1:**
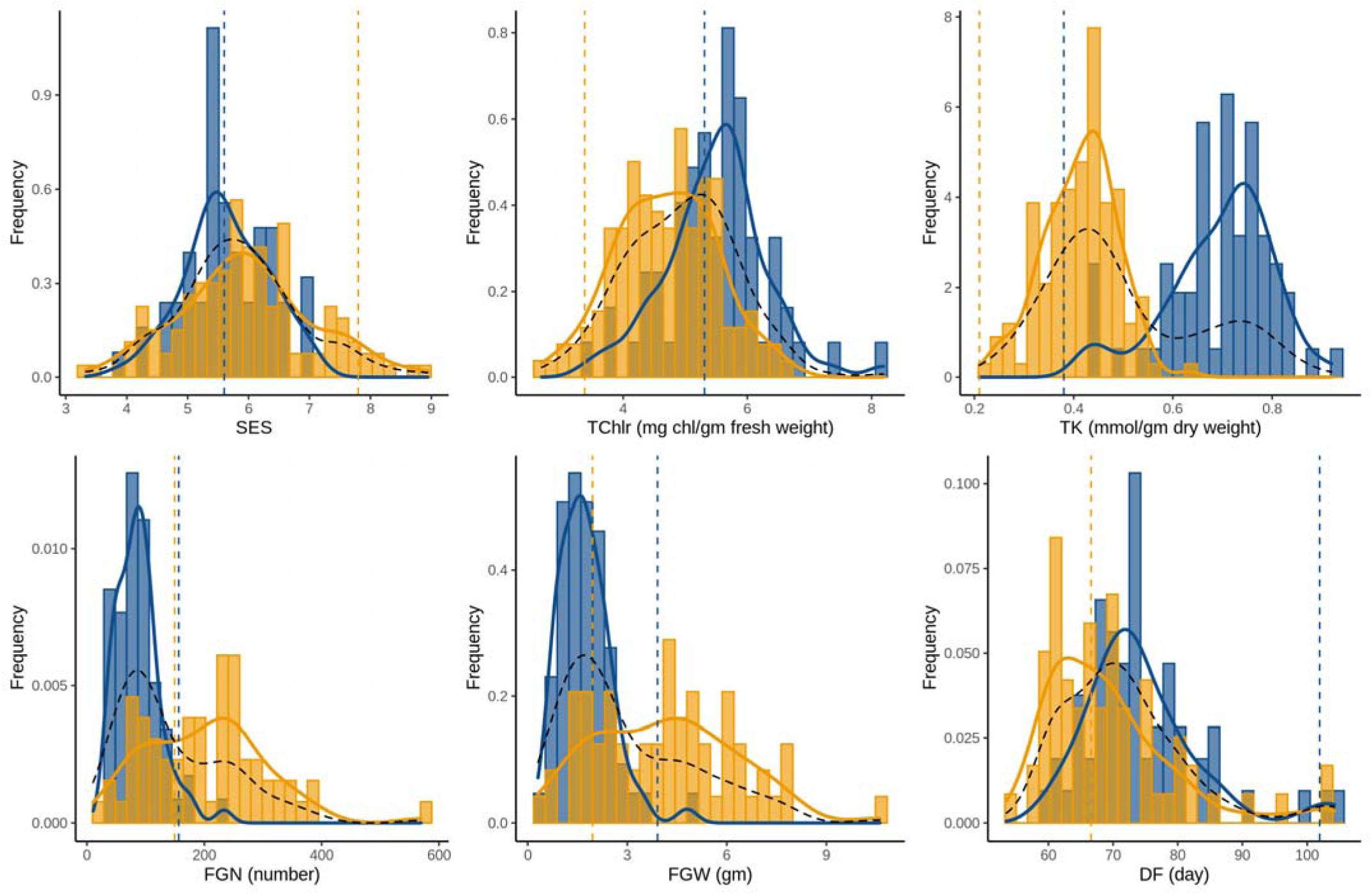
Frequency distribution of traits showing transgressive segregation in the F_2_ population and the individual subsets of cross directions. Blue and orange histograms indicate samples from *Horkuch*♀ and *IR29*♀ cytoplasm respectively. Curves in blue and orange indicate distribution plots of *Horkuch*♀ and *IR29*♀ cytoplasm respectively and dotted curve in black indicates the distribution plot of total population. Parental values are marked by a dotted vertical line where blue indicates *Horkuch* and orange indicates *IR29*.

To understand the partitioning of genetic variation we used principal component analysis of 130 families which had complete observations of all traits for both the stages of treatment. The first two PC axes comprise the majority of genetic trait variation in this population (31.3%) (Figure 2). This indicated substantial genetic correlation among these traits in this population. Interestingly, principal component one separates individuals into two groups almost exclusively depending on their respective cytoplasm. This observation further supports a significant role of cytoplasm on the performance of a plant under different treatment stages. We also found moderate correlation of some traits between the two different stages of treatment: PH showed positive correlation with SL and yield related traits such as FGN, FGW, SF, HI and PE is negatively correlated with TK (Figure 2). This correlation suggests some possible shared mechanisms of salinity responses that has some trade-off between two different treatment stages.

**Figure 2:**
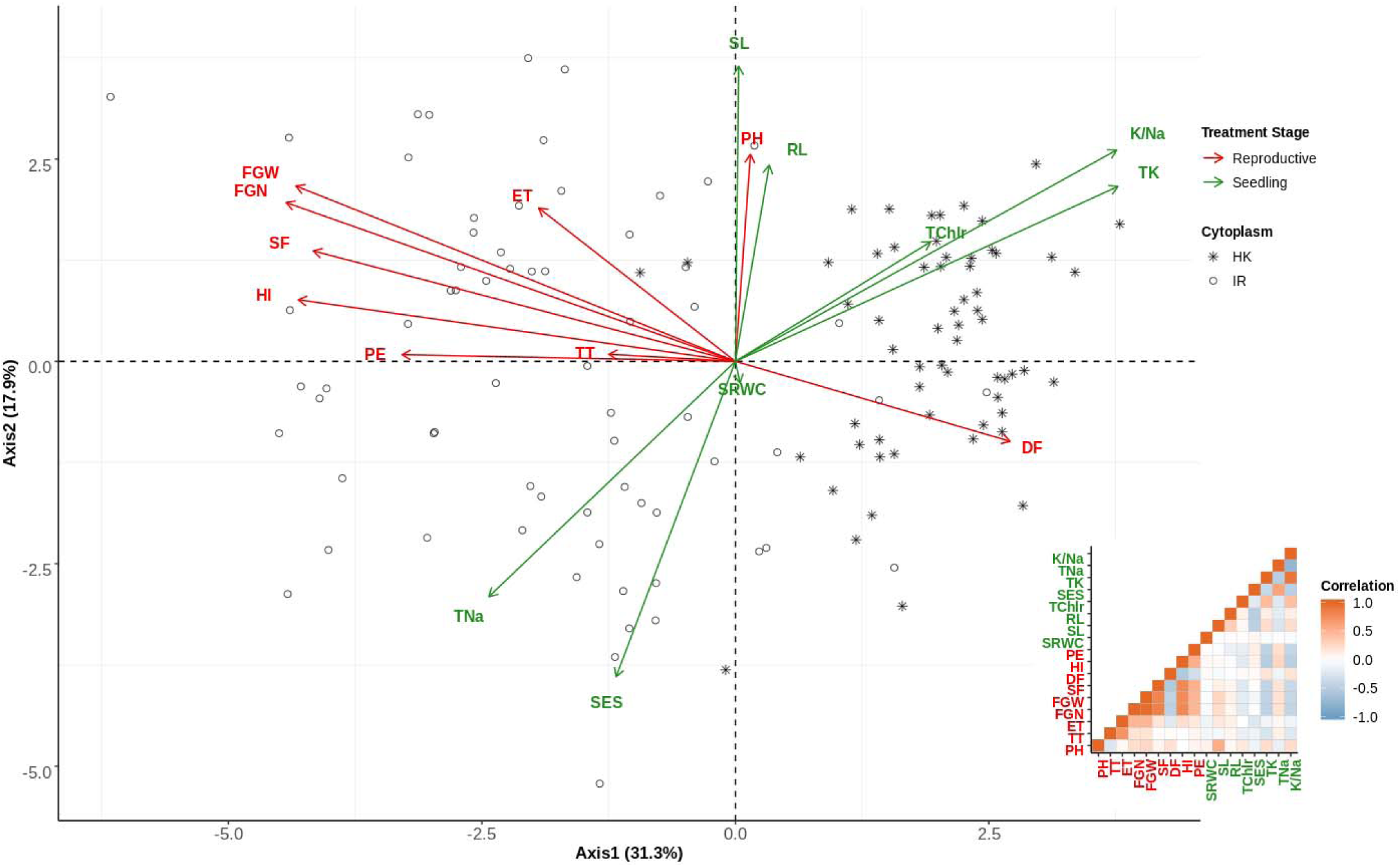
PCA on trait correlations in the F_2_ mapping population. Each point represents the genetic means of each F_2_ family whereas the shape of point indicates the cytoplasm (cross direction). Direction of variation for axis 1 and 2 of each trait has been plotted as arrow and are color labelled depending on two different treatment stages: green indicates Seedling stage treatment and red indicates reproductive stage treatment. Labels of traits are printed close to the arrow-head. Small insert-plot at the bottom-right shows the correlations of traits where brown color shows positive correlation and light-blue indicates negative correlation. Traits labelled with green color indicates seedling stage ones and red indicates reproductive stage traits.

### Linkage map construction

In our previous study we constructed a linkage map for this reciprocal mapping population by applying ddRAD technique (Noor *et al*., 2019). Unfortunately, we had high genotyping error and inflated segregation distortion for the two parental alleles for a given locus. Therefore, we failed to capture the genome-wide linkage map which resulted in the removal of the entire chromosome 5 for QTL mapping. In this study, we genotyped this reciprocal mapping population using DArTSeq (Diversity Array Technology coupled with Sequencing) technique in order to improve genotyping error by adding higher sequence coverage. DArTSeq can generate low to moderate density SNP information with high coverage and low cost. This method uses restriction enzymes to reduce the complexity of the genome and has been optimized for various plant species to achieve best complexity reduction. We used this platform for our mapping population to generate genotype information at ∼10 thousand loci that are well-distributed in the rice genome. For analyses, we filtered all the DArTseq SNP loci to obtain polymorphic homozygous SNPs for the parents and retained only loci that had minimum 50% representation in the population. With these filter criteria we obtained 2,230 high quality SNPs for this mapping population. Among these loci, 956 markers that showed significant segregation distortion by chi-square test [P value < 1e-2] were removed. More distorted loci were skewed towards the *Horkuch* parent than the *IR29* parent. 739 markers that were similar were dropped using a minimum recombination fraction threshold of 0.03. Markers were reordered by the concorde program as determined by the tspOrder function (Monroe *et al*., 2017). The final map was constructed by “kosambi” map function after dropping 36 markers which had very different orders between their genetic map and physical position in genome. The final map had 499 markers with a map size of 2004.8 cM and average distance between markers of 4.1 cM (Supplementary Figure 1A). The maximum gap of 22.3 cM was found in chromosome 5. Chromosome 7 had the fewest markers (22 markers). Supplementary Figure 1B presents the concordance of genetic map with the physical map of rice genome. As mentioned in earlier, we detected significant association of cytoplasm for multiple traits in both stages of treatment therefore we aimed to test (Chi-square test of independence, see Method section) for cytoplasm-nuclear association for each marker in this genetic map. We detected 10 loci that showed significant cytoplasm-nuclear association [FDR < 0.1] and these are mostly clustered in chromosomes 2, 3 and 7. This association suggested the presence of cytoplasm-nuclear linkage disequilibrium in this mapping population resulting from selection at the reproductive stage during the development of the mapping population. Overall, we constructed a linkage map with moderate marker density that was closely aligned with the physical map of rice genome.

### Significant QTL at both growth stages QTL for seedling stage treatment

We measured eight traits that reflect the survival performance of rice seedlings under salinity stress. We found six QTL for three traits: SL, RL and TK (Table 1, Figure 3). We found three significant QTL for SL occurring at qSL.1@183 (reporting a QTL for SL located at chromosome 1 at 183 cM), qSL.3@218, and qSL.5@160. For these QTL, the positive alleles were from *Horkuch* parent and we detected no significant cytoplasmic effects or interactions. The QTL at qSL.1@183 had a large effect corresponding to a ∼3.5 cm increase in seedling length that has a confidence interval of 3.5 Mbp and localized at the end of chromosome 1 (∼38.4 Mbp). The second QTL, qSL.5@160, had a small confidence interval of ∼ 1 Mbp but the effect size was moderate. The third QTL at qSL.5@160 had a very larger confidence interval (∼ 9 Mbp) with small effect size.

**Figure 3:**
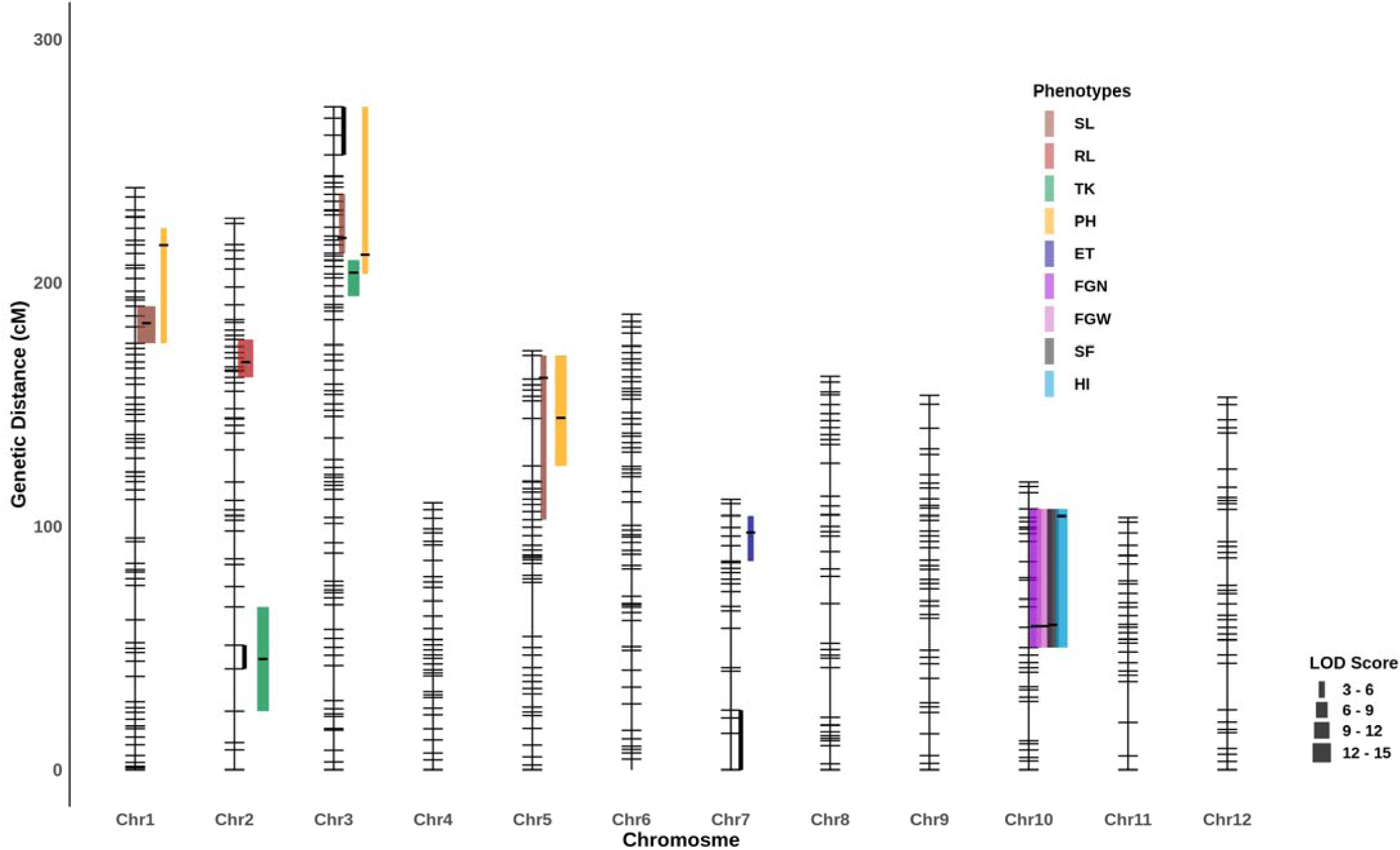
Illustration of QTL across chromosomes. QTL are denoted as a point and 1.5 LOD drop confidence intervals extended to a true marker is indicted by the bar for each QTL. Peaks of the QTL were marked as black line in QTL intervals. QTL from same trait are marked with same color. Line width represents the magnitude of LOD score. Genomic regions that showed significant association with cytoplasm are marked here with black line segment.

We identified one large effect QTL for RL at qRL.2@167 with a confidence interval of ∼ 4 Mb. Here, the *Horkuch* parent contributed the positive allele. For TK, cytoplasm was a significant covariate that interacted with two QTL which were detected at qTK.2@45 and qTK.3@203. In addition, the QTL qTK.2@45 is co-localized with the cluster of genetic loci that showed significant cytoplasm-nuclear association. The other QTL for TK, qTK.3@204 did not overlap with the association cluster in chromosome 3 but resided in close proximity. This evidence suggests a possible role of cytoplasmic-nuclear interaction for this trait.

### QTL for reproductive stage treatment

Very little work has been published on salinity stress during rice reproductive stages. For this stage, our primary focus was on yield responses of the plants under salt stress. We found 8 QTL for 6 traits at reproductive stage salinity treatment (Table 1, Figure 3). A major effect QTL for FGN was found on Chromosome 10 at 58.5 cM (qFGN.10@58.5). Here, the *IR29* parent contributed the positive allele. We found significant cytoplasm-nuclear interaction for QTL model where the *IR29* allele had positive effects only for *IR29*♀ (Figure 4). However, this QTL had a very wide confidence interval of ∼6 Mbp. We found two co-localized QTL in this same region including a QTL for FGW and another for SF at <u>qFGW.10@58.5</u> and qSF.10@59 respectively. These two QTL also had significant interactions with cytoplasm in their corresponding QTL models. *IR29* contributed the positive allele for both <u>qFGN.10@58.5</u> and <u>qFGW.10@58.5</u>. For FGW, as like FGN, the *IR29* allele had a positive effect only for *IR29*♀ but for SF this allele not only had positive effect for *IR29*♀ but also had a negative effect for *Horkuch*♀. We also found a QTL for HI at qHI.10@104 for which the positive allele was from IR29. However, this model only had an additive effect of cytoplasm.

**Figure 4:**
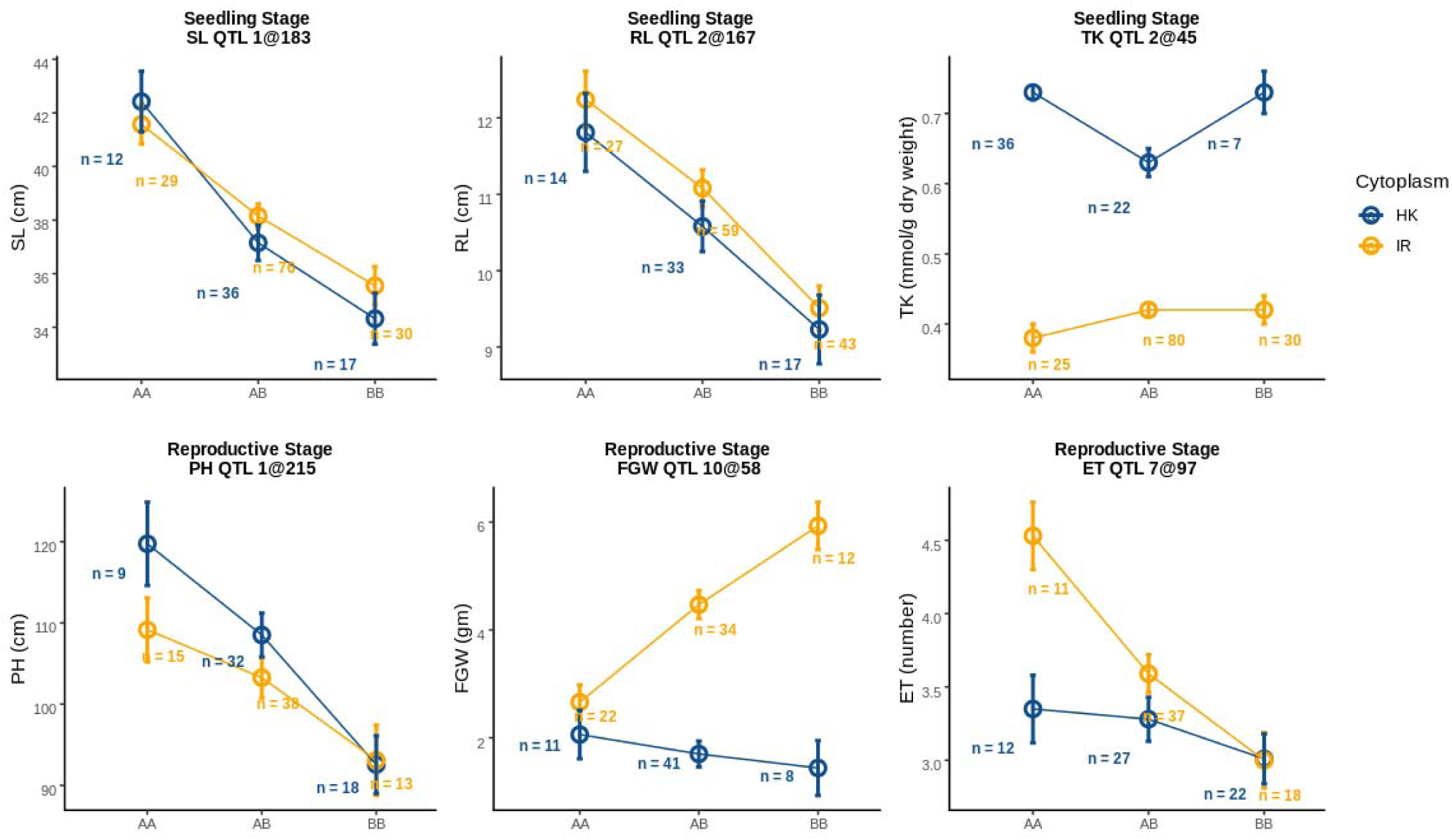
Interaction plots of allelic effect of QTL and cytoplasm on different traits from two different treatment stages. Blue line shows plants with *Horkuch* cytoplasm whereas orange line indicates plants with *IR29* cytoplasm. Alleles are plotted on x-axis where AA, AB and BB indicate homozygous *Horkuch*, heterozygous of *Horkuch*/*IR29* and homozygous *IR29* respectively. Allelic means +/- SE are reported. Representative QTL effects for SL and PH are presented in the upper panel and exhibit no significant interaction with cytoplasm. The third plot from the left on upper panel demonstrates significant additive effects of the maternal cytoplasm on TK. In the bottom panel, plot two and three from the left demonstrate significant interaction of QTL alleles with cytoplasm for traits such FGW, ET.

Another QTL, qET.7@97 cM was detected for ET where the *Horkuch* parent contributed the positive allele. Similar with the other grain related traits, cytoplasm contributed significantly to this QTL model. However, the positive allele from *Horkuch* performed better in *IR29*♀. We also found three PH QTL occurring at qPH.1@215, qPH.3@211 and qPH.5@144, for which the positive alleles were from the *Horkuch* parent. The first two QTL models showed only additive contribution of cytoplasm but the third showed also an interactive effect of cytoplasm. Overall, we found a hotspot of QTL on chromosome 10 for multiple parameters related to yield. The significant correlation of these traits may reflect related metrics of yield performance in rice.

### QTL co-localization

In this study, we found that some QTL intervals of various traits overlapped, therefore we annotated these overlapping intervals as QTL clusters. We detected a co-localized QTL at 1@175:220 cM [QTL Cluster 1 (QC 1)] affecting PH and SL from two different stages of treatment. These traits represent the vigor of plants at the two different growth stages and had significant positive correlation (Pearson’s correlation coefficient = 0.54). In parallel with this co-relation, these two QTL had positive alleles from the *Horkuch* parent which indicate the same sign of effects for these QTL. Another wide co-localized cluster was found at 3@194:273 cM [QTL Cluster 2 (QC 2)] impacting SL, PH and TK with the positive allele from the *Horkuch* parent. However, TK showed no significant correlation with PH which was causal for the co-localization for QC1. A third QTL Cluster 3 (QC 3) was found at 5@144:170 for SL and PH where the positive alleles were from *Horkuch* parent. The fourth QTL Cluster (QC 4) at 10@58:107 was found for four yield related traits including FGN, FGW, SF and HI for which the positive alleles were from the *IR29* parent.

### GO enrichment analysis of candidate genes within QTL confidence intervals

To understand the molecular mechanism of salt tolerance, we further investigated the function of candidate genes that were located within the QTL confidence intervals (Supplementary File 1). We applied GO enrichment analysis on the candidate gene lists against the genome-wide background frequency of GOs. The results for GO enrichment analysis have been provided in Supplementary File 2. For the seedling stage, QTL at qSL.1@183 showed significant enrichment of GO terms such as pollination, protein lipidation, lipoprotein and liposaccharide metabolic process, various transport and DNA-directed RNA polymerase complexes. Another QTL for SL at qSL.3@218 was significantly enriched with the GO terms protein transport and localization, amide and lipid transport, cytoskeleton and actin binding. The third QTL for SL, qSL.5@160 had significant enriched GO terms for chromatin assembly, nucleosome organization, DNA packaging, ion transport, amide and peptide transport. QTL qRL.2@167 showed significant enrichment of GO terms such as carboxylic, dicarboxylic and C4-dicarboxylate transport, malate transport, anion transport and sexual reproduction. Two QTL for TK, qTK2@45 and qTK3@204, were both enriched with the GO terms oxidoreductase activity, various transmembrane transporter activity, potassium ion transmembrane transporter and cation transmembrane transporter.

For traits at reproductive stage, the QTL qPH.1@215 was significantly enriched with the GO terms anion and potassium ion transmembrane transport, divalent metal ion transport, pollination, reproduction process, endoplasmic reticulum and organelle sub-compartment. Another PH QTL, qPH.3@211 showed enrichment for GO terms such as cellular nitrogen compound metabolism, organic acid transport, mitochondrial membrane and protein complex. The third QTL, qPH.5@144 also showed significant enrichment for nitrogen compound metabolic process. This QTL was also enriched for various mitochondria and cytoplasm related GO terms. The QTL, qET.7@97 was enriched with cytoplasm and mitochondrial membrane part and chlorophyll metabolic process. QTL for FGW, FGN, SF and HI share the same QTL intervals therefore the gene models within this interval were identical. This interval was enriched with cell wall macromolecule catabolic process, amino sugar and glycan metabolic process, protein localization to organelles and mitochondrial transport. The significant enrichment of mitochondrial and organelle related GO terms for some QTL confidence intervals suggests a possible explanation for the significant cytoplasm and cytoplasmic-nuclear interactions detected in our study.

For QTL models that had significant cytoplasmic effect, 1473 annotated genes are present in the respective confidence intervals. In our previous studies we tested for the association of gene expression to salinity stress for this same reciprocal mapping population at seedling and reproductive developmental stages (Razzaque *et al*., 2019; Razzaque *et al*., 2017). Among the genes that are present in the QTL confidence interval, 188 showed significant cytoplasm*treatment interaction in these previous gene expression studies. In order to identify their association for cellular component we further tested for enrichment of GOs for these common genes that are present in QTL intervals and showed significant cytoplasm*treatment interaction in our previous gene expression studies. (Supplementary File 3). We found these genes to be significantly enriched with GO terms such as mitochondrial proton-transporting ATP-synthesis complex, mitochondrial protein-complex and mitochondrial membrane.

## DISCUSSION

In this study, we explored the responses of rice to salinity stress at two different growth stages using a reciprocal mapping population. Among the 14 QTL that we reported, 8 QTL models showed significant effect of cytoplasm. This finding underlines the importance of considering both organelle and nuclear genome for complex traits such as salinity tolerance.

Cytoplasmic background may play an important role in trait genetic architecture by itself or through complex interactions with the nuclear genome. (Joseph *et al*., 2013b; Lovell *et al*., 2015; Moison *et al*., 2010; Tang *et al*., 2013). Gregorio and Senadhira (1993) reported significant reciprocal effects among crosses for salinity response in their study of nine different rice varieties and suggested using susceptible plants as male parent for hybridization programs. In order to identify the best candidates for QTL pyramiding by breeders, it is essential to estimate single QTL effect for the trait of interest. Hence, it is important to test for the random effect of covariates such as cytoplasm in a QTL model and estimate its effect size. In this way, the causality, contribution and combinations of cytoplasm and nuclear-donor alleles of QTL can be defined. Moreover, including cytoplasm as covariate in QTL mapping can increase the ability to detect small-effect QTL peaks if there is a significant contribution of cytoplasm for that given trait. Considering all these aspects, we employed a QTL modeling framework where the cytoplasm-nuclear interaction was also considered as a contributor to phenotypic variance. For TK, FGN, FGW the additive effect of cytoplasm was significantly large compared to the effect of a single QTL (Figure 4). Overall, we found significant contribution of cytoplasm for traits related to yield, such as FGN, FGW and SF as well as one important trait for the seedling stage TK. Identification of causal impacts of cytoplasm will help to define the best combination of cytoplasm and nuclear-donor materials and will underscore the selection trade-off for multiple desired traits. For instance, on the one hand, we found the positive nuclear allele of *IR29* had its effect only in *IR29*♀ for the QTL model of yield related traits; while on the other hand we observed the strong positive effect of *Horkuch* cytoplasm for the QTL model of TK at the seedling stage treatment. The latter trait is a highly desired one for breeding of salt tolerant varieties. Hence, estimating the contributions of cytoplasm for multiple traits can help understand the performance trade-off in a breeding program for QTL pyramiding.

The cytoplasmic genome can influence interaction of alleles from nucleus and cytoplasm and can favor the evolutionary co-adaptation of high-fitness. In the current study we found a significant association of cytoplasm for some traits and therefore further tested for non-random interaction of alleles for nucleus and cytoplasm. We found that the QTL qTK.2@45 was a hotspot of cytoplasm-nuclear interaction on chromosome 2. Additionally, qPH.5@144 was another similar hotspot on chromosome 5. Both of these QTL models showed significant effect for cytoplasm. For qTK2@45, the effect of cytoplasm was mostly additive where *Horkuch*♀ contributed large positive effect. On the contrary, for qPH5@144, cytoplasm had an interactive effect. *IR29* nuclear allele had a negative effect on PH and homozygous *IR29* nuclear allele on *Horkuch*♀ had even larger negative effect size (Supplementary Figure 3). Taken together, this suggests a significant interaction of nuclear alleles with the cytoplasmic genome. This further supports the fact that selection of the female plant plays an important role for the performance of a breeding population and while pyramiding QTL and the conditional selection of cytoplasm may have some trade-off on a hybrid plant’s performance. Moreover, we detected significant cytoplasm-nuclear linkage of a few markers that overlapped with some QTL intervals. Therefore, careful consideration is needed in order to select these loci for QTL pyramiding.

One important finding in this study is that we have detected multiple co-localized QTL within and among the two different stages of salinity treatment. This finding emphasizes the possible constraints during selection of QTL in a breeding program. Here we identified four QTL clusters where multiple trait QTL co-localized. Co-localized QTL can impose constraints on selection for QTL pyramiding. As an example, we found that QTL for PH and SL are colocalized at QTL cluster 1 on chromosome 1. This cluster had a positive effect for the *Horkuch* parental allele for PH and SL. However, a taller plant is not the desired plant architecture for a breeding program for high-yielding rice varieties since this will lead to over-investment of energy in vegetative growth and potential lodging. On the other hand, Leon et al. (2015) reported that percent of shoot length reduction under saline treatment is highly co-related to saline sensitivity. This conditional relationship between traits results in some possible trade-offs between favorable and undesirable traits. The same logic is applicable for QTL cluster 2 where traits (SL, TK and PH) for these clusters are positively correlated but increased PH is not desirable for any breeding program. On the other hand, for QTL cluster 4, all the yield related such as FGN, FGW, SF and HI could be combined where the *IR29* parent contributes all the positive alleles. Taken together, these findings underscore the importance of studying the performance of a plant for different developmental stages. In addition to that, we need to consider the fact that selection for multiple traits may not be orthogonal due to the complex mechanisms of salt adaptation.

To understand the molecular mechanism of salt response and the effect of cytoplasm for salt tolerance we tested for enrichment of GO functions for genes within QTL confidence intervals. Both the QTL intervals for TK were enriched with various transmembrane transporter activity, and potassium ion transmembrane transporters. K^+^ is involved in numerous metabolic processes in plants and excess Na^+^ interferes with the K^+^ homeostasis during salinity stress. To maintain the cellular homeostasis of K^+^, various potassium transmembrane transporters have been reported to increase salt tolerance in various glycophytes (Tester and Davenport, 2003). In the qTK3@204, a specific peroxidase was detected as a cis expression QTL (Seraj et al, Unpublished data). Peroxidases normally reduce reactive oxygen species (ROX) under stress and can contribute to regulation of HAK type potassium transporters. Also, in the qTK2@45 region a calcium transporting ATPase was detected as a cis eQTL that showed differential expression in the two parents (Seraj et al, unpublished data). This is a less characterized plasma membrane calcium ATPase in rice named OsACA2 (Singh *et al*., 2014) that catalyzes the hydrolysis of ATP coupled with the transport of calcium in cytosol and maintains calcium homeostasis under salt stress. It shows close similarity with OsACA5 (LOC_Os04g51610), in some studies known as OsACA6, which has been previously characterized to enhance salt stress tolerance (Huda *et al*., 2013). Normally under salt stress an increase in Ca^2+^ ensues in a time-sensitive manner and the homeostasis is maintained by Ca^2+^ channels, Ca^2+^ exchangers and Ca^2+^ ATPases. Also, Calcium signaling pathway has an important role in activation of potassium channels needed to maintain the potassium homeostasis under salt stress. Role of this differentially expressed calcium ATPase in K^+^ QTL region along with other calcium signaling components and potassium channels/transporters needs careful characterization.

For the reproductive stage, we found most of the QTL intervals for PH, ET, FGW, FGN, SF and HI were enriched with mitochondria, cytoplasm and organelle related GOs. This supported the observation that these QTL models also showed significant interaction with cytoplasm. Thus, a possible interaction of the cytoplasm genome with nuclear alleles present in the region of QTL confidence intervals is likely. Additionally, enrichment analysis of DEGs (significant for cytoplasm*treatment model) from our previous studies (Razzaque *et al*., 2019; Razzaque *et al*., 2017) on this mapping population were enriched with GO terms such as organelle, thylakoid, mitochondria, photosynthesis, cation transmembrane transporter and various sodium symporter activities. Salt stress inhibits photosynthesis of plants but how this affects the ionic balance of chloroplasts has not been studied much, until recently. Bose et al. (2017) has proposed some candidate transporters that are involved for the movement of sodium, potassium and chloride across chloroplast membrane in glycophytes and halophytes and explained how these transporters may regulate photosynthesis in chloroplast. These candidate symporters include bile acid: sodium symporter and cation transmembrane transporter which have possible role in maintaining chloroplast ion homeostasis. From our gene expression studies of cytoplasm*treatment DEGs, enrichment of symporter GOs that are localized in mitochondria and organelles suggest a possible role of mitochondria and chloroplast during salinity stress and tolerance or sensitivity to it. The bile acid: sodium symporter gene also appeared as a trans expression QTL under salt stress linked with the potassium QTL region qTK2@45 as well as <u>qPH.3@211</u>. (Seraj et al, unpublished data). This evidence also suggests a plausible explanation why we found cytoplasm as a covariate in QTL models for this study. These are likely candidates for future functional genomic studies of salinity tolerance in rice.

In this QTL analysis framework, we applied linear mixed model which can handle cytoplasm and alleles as fixed effect predictors. This model can also consider residual polygenic variation as a random effect using a kinship matrix. Here we implemented DArtSeq technique which can genotype a moderate number of SNPs that are well-dispersed in rice genome and aimed to select SNPs close to gene space of the rice genome. In our previous study, we had generated a genetic map on this mapping population by ddRAD technique which failed to capture a significant space of genetic map due to erroneous genotyping and high rate of missing SNP calls (Noor *et al*., 2019). In this current study, we implemented a robust QTL analysis framework on this improved genetic map and we were able to detect three QTLs for SL and one RL at seedling stage salinity treatment which we could not detect in our earlier study (Supplementary Table 2; Supplementary Figure 4). For reproductive stage salinity treatment, we were able to detect additional five QTL for PH, ET and SF. We have also detected one big effect QTL for FGN and FGW in a different chromosome in this current study due to the fact that in our previous study we failed to capture markers at that region. In addition to that, this framework provided QTL with higher likelihood and tighter confidence interval and provided better estimation of effect size of each QTL for a given trait. Therefore, the additional detected QTL with high LOD scores and tighter confidence intervals may contribute significantly for the improvement in development of rice which combines both salt tolerance and high yields.

## CONCLUSION

In this study, we aimed to identify genetic loci for salinity tolerance of a rice landrace, *Horkuch*, at two sensitive developmental stages. We found 14 QTL for 9 traits under salinity treatment. We detected some overlap in the genomic regions affecting traits across developmental stages. One chief finding of this study was the significant contribution of cytoplasm on many traits and eventually its effect on the corresponding QTL model. Enrichment analyses suggest that the observed cytoplasmic effect could be causally related to plastid symporter activity and their interaction with nuclear genes. Collectively, this study helped to understand the genetic basis of salt tolerant mechanism of a local rice landrace *Horkuch*. Moreover, careful implementation of pyramiding of QTLs that were detected in this study can pave a way to generate high yielding salt tolerant rice varieties.

## Supporting information

Detailed supplementary methods, tables and figures

List of genes in QTL confidence interval where one sheet represents one single QTL

List of significant GO terms for genes in QTL confidence interval where one sheet represents one single QTL

List of significant GO terms for the common genes between all candidate genes of QTL confidence interval and significant DEGs for cytoplasm*treatment

## SUPPLEMENTARY DATA

**Supplementary Material:** Detailed supplementary methods, tables and figures

**Supplementary_File_1:** List of genes in QTL confidence interval where one sheet represents one single QTL

**Supplementary_File_2:** List of significant GO terms for genes in QTL confidence interval where one sheet represents one single QTL

**Supplementary_File_3:** List of significant GO terms for the common genes between all candidate genes of QTL confidence interval and significant DEGs for cytoplasm*treatment interaction model of seedling shoot tissue, seedling root tissue and reproductive shoot tissue

## CONTRIBUTIONS

Z.I.S., S.M.E. and M.S.R. designed the experiment. M.S.R. did the reciprocal crossing. S.M.E., S.R., S.F.K., S.B. and G.M.N.A.J grew the plants and collected the phenotypes. S.M.E. and T.H. analyzed phenotypes. S.M.E. and S.R. isolated DNA for DArTseq. T.H and S.M.E did genotype calling from raw data. T. H. did the modeling for QTL and other statistical analyses. T.E.J provided feedback on the statistical models and analyses. T.H. wrote the manuscript. T.E.J. and Z.I.S. provided their significant feedback for writing. S.M.E. and S.R. revised the manuscript.

## ACKNOWLEDGEMENT

Thanks to Nazrul Islam and other technicians at BRRI and Dhaka University who took care of the plant materials. We would like to thank the members of the Plant Biotechnology Lab and Juenger Lab who provided their feedback on this research project. This work was supported by the PEER science grant to Z.I.S. by the National Science Foundation (Project 2-004) and USAID (agreement number AID-OAA-A-11-00012). The grant was hosted by an existing NSF grant from the Plant Genome Research Program (IOS-0922457) to T.E.J. S.M.E. was supported through a PhD fellowship grant from Monsanto’s Beachell-Borlaug International Scholars Program (MBBISP). We would like to thank Dr. Harkamal Walia for supervising S.M.E for her MBBISP. We are thankful to Dr. Adam Price for recommending DArtseq to us for genotyping and also for supervising S.F.K. in her Split Commonwealth Fellowship. We would also like to thank Dr. Abdelbagi M. Ismail to allow us to use IRRI facilities for raising the reciprocal population.

