## Supplementary material for "Natural variation in growth and physiology under salt stress in rice: QTL mapping in a *Horkuch* × *IR29* mapping population at seedling and reproductive stages": Detailed supplementary methods, tables and figures

### 1 Supplementary Methods

**1.1	Comparison of QTL models between new and old linkage maps**

The first linkage map of this bi-directional cross had constructed from ddRAD genotyping data using R/qtl package (Noor *et al.*, 2019).Unfortunately, we found very high segregation distortion for genetic markers and were failed to capture a substantial amount of genetic space including whole chromosome 5. In this study we used DArtSeq technique to generate moderate number of SNP that were evenly distributed across rice genome and were aimed to capture genetic space close to gene space. In order to compare different QTL models, build by using two different genetic maps, we plotted the physical position of each true marker on x-axis and LOD score for each model on y-axis (Supplementary Figure 4). We excluded pseudo-markers while plotting LOD profiles for each QTL model.

### 2 Supplementary Figures and Tables

#### 2.1 Supplementary Tables

|  | Phenotypes | Abbreviation | Population | | Parent | |
| --- | --- | --- | --- | --- | --- | --- |
|  |  |  | Mean | SD | *Horkuch* | *IR29* |
| Seedling Stage | Shoot Relative Water Content (%) | SRWC | 70.06 | 5.07 | 67.72 | 62.43 |
|  | Standard Evaluation System (number score) | SES | 5.83 | 0.99 | **5.60** | **7.80** |
|  | Shoot Length (cm) | SL | 38.09 | 4.64 | **41.78** | **21.35** |
|  | Root Length (cm) | RL | 10.71 | 2.15 | **11.51** | **7.60** |
|  | Total Chlorophyll ***(mg chl per gram fresh weight) | Tchlr | 5.04 | 0.92 | **5.31** | **3.38** |
|  | Total Sodium *** (mmol/g dry wt) | TNa | 3.07 | 0.77 | **3.08** | **6.23** |
|  | Total Potassium *** (mmol/g dry wt) | TK | 0.38 | 0.08 | **0.38** | **0.21** |
|  | Potassium by Sodium (ration) ** | K/Na | 0.14 | 0.04 | **0.13** | **0.04** |
| Reproductive Stage | Plant Height (cm) | PH | 104.5 | 17.61 | **129.81** | **75.50** |
|  | Panicle Exsertion (%) *** | PE | 99.24 | 1.42 | 100.00 | 98.81 |
|  | Total Tiller*** (number) | TT | 4.66 | 1.16 | 5.07 | 4.36 |
|  | Effective Tiller (number) | ET | 3.43 | 0.8 | **3.87** | **3.23** |
|  | Filled Grain Number (number) *** | FGN | 140.87 | 92.96 | 156.29 | 149.85 |
|  | Filled Grain Weight (gm) *** | FGW | 2.85 | 1.95 | **3.91** | **2.00** |
|  | Spikelet Fertility (%) *** | SF | 48.63 | 15.93 | **49.4** | **45.08** |
|  | Days to Flower (day) *** | DF | 72.08 | 9.83 | **101.93** | **66.60** |
|  | Harvest Index***(ratio) | HI | 0.26 | 0.12 | 0.23 | 0.32 |

**Supplementary Table1:** Descriptive statistics of phenotypes measured at seedling and reproductive stages under salinity stress.

Asterisks indicates significant effect of cytoplasm (*** for P < 0.001, ** for P <0.01 and * for P <0.05). For the parent’s column, trait values in bold character indicate significant difference between the group means of two parents (p-value < 0.05)

**Supplementary Table 2:** Comparison of detected QTL in current study vs. previous study

|  | Number of QTL | QTL names |
| --- | --- | --- |
| Newly detected QTL in current study | 9 | qSL.1@183  qSL.3@218  qSL.5@160  qRL.2@167  qPH.1@215  qPH.3@211  qPH.5@144  qET.7@97  qSF.10@59 |
| QTL that changes chromosomal location | 5 | qTK.2@45  qTK.3@204  qFGN.10@58  qFGW.10@58  qHI.10@104 |

#### 2.2 Supplementary Figures

**Supplementary Figure 1.** A. Genetic map of this F_2_ mapping population where each vertical line segment represents one chromosome and horizontal line segment of each chromosome represent the position of genetic marker in CM. B. Plot of physical vs genetic distance of markers for the linkage map.

**Supplementary Figure 2.** Interaction plots of allelic effect of QTL and cytoplasm at seedling stage salinity treatment. Blue line shows plants with *Horkuch* cytoplasm whereas orange line indicates plants with *IR29* cytoplasm. Alleles are plotted on x-axis where AA, AB and BB indicate homozygous *Horkuch*, heterozygous of *Horkuch*/*IR29* and homozygous *IR29* respectively. Allelic means +/- SE are reported.

**Supplementary Figure 3.** Interaction plots of allelic effect of QTL and cytoplasm at reproductive stage salinity treatment. Blue line shows plants with *Horkuch* cytoplasm whereas orange line indicates plants with *IR29* cytoplasm. Alleles are plotted on x-axis where AA, AB and BB indicate homozygous *Horkuch*, heterozygous of *Horkuch*/*IR29* and homozygous *IR29* respectively. Allelic means +/- SE are reported.

**Supplementary Figure 4:** Comparison of two different QTL models for TK and FWG. Estimated LOD score for each model were plotted against the corresponding physical position of true markers. Orange points represent the LOD profile for new linkage map built using DArtSeq technique while blue points represent the old linkage map constructed from ddRAD data. Chromosomes that had one significant QTL in either model have been plotted.

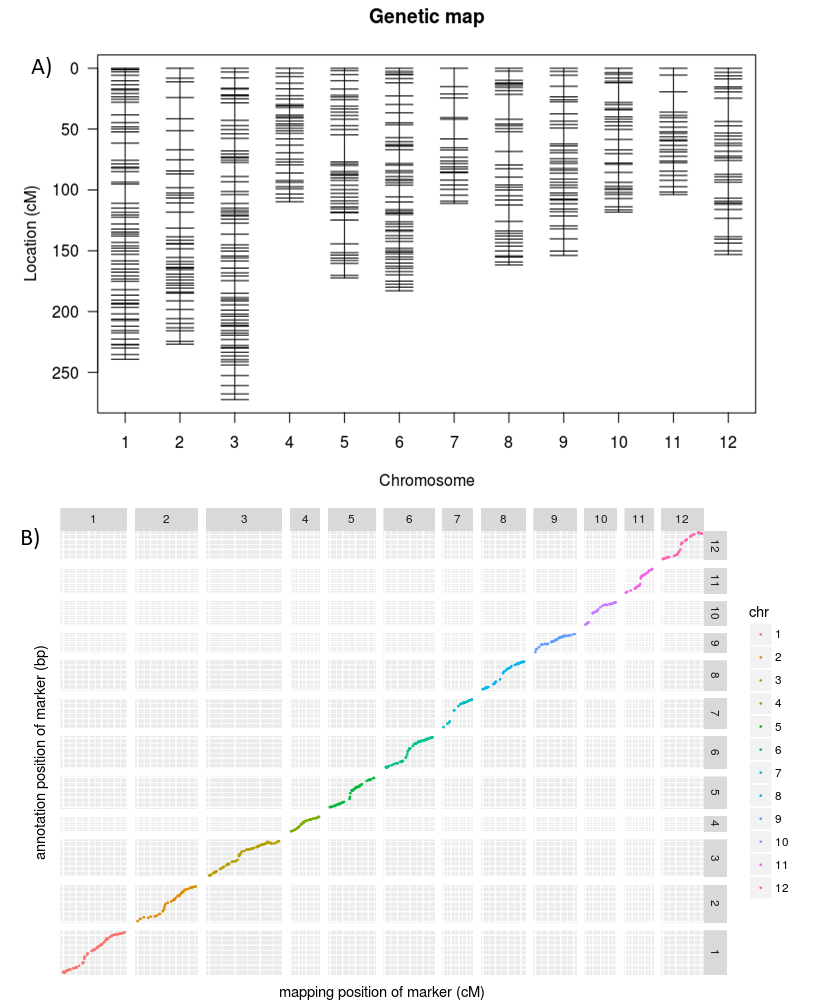

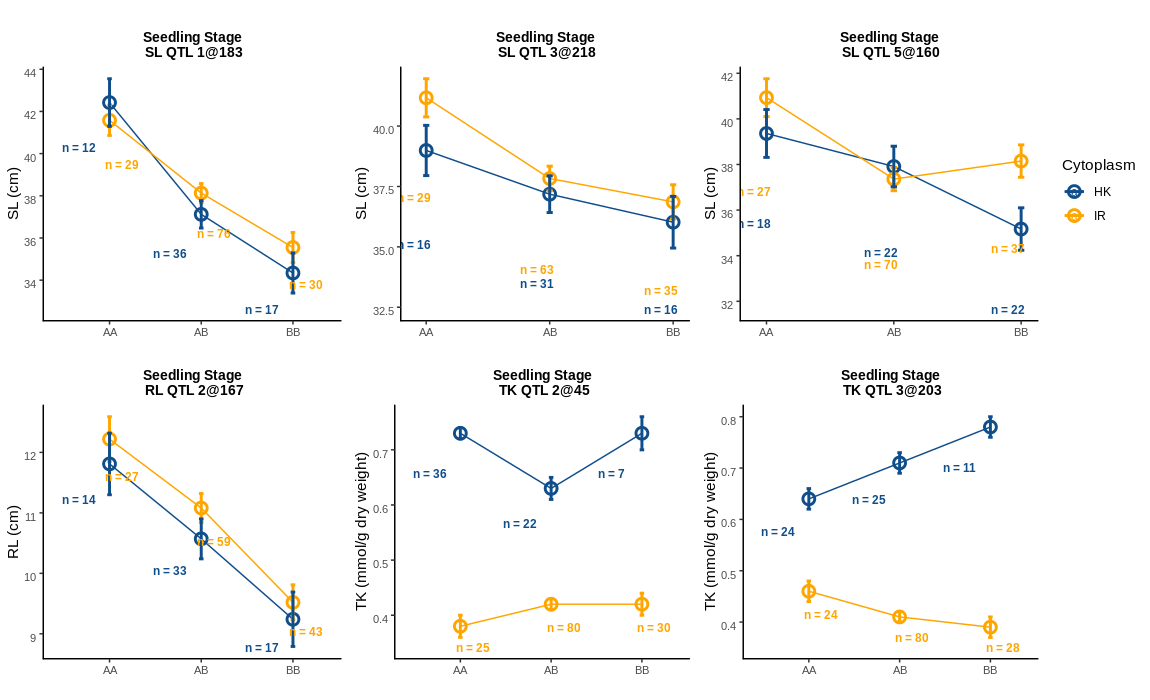

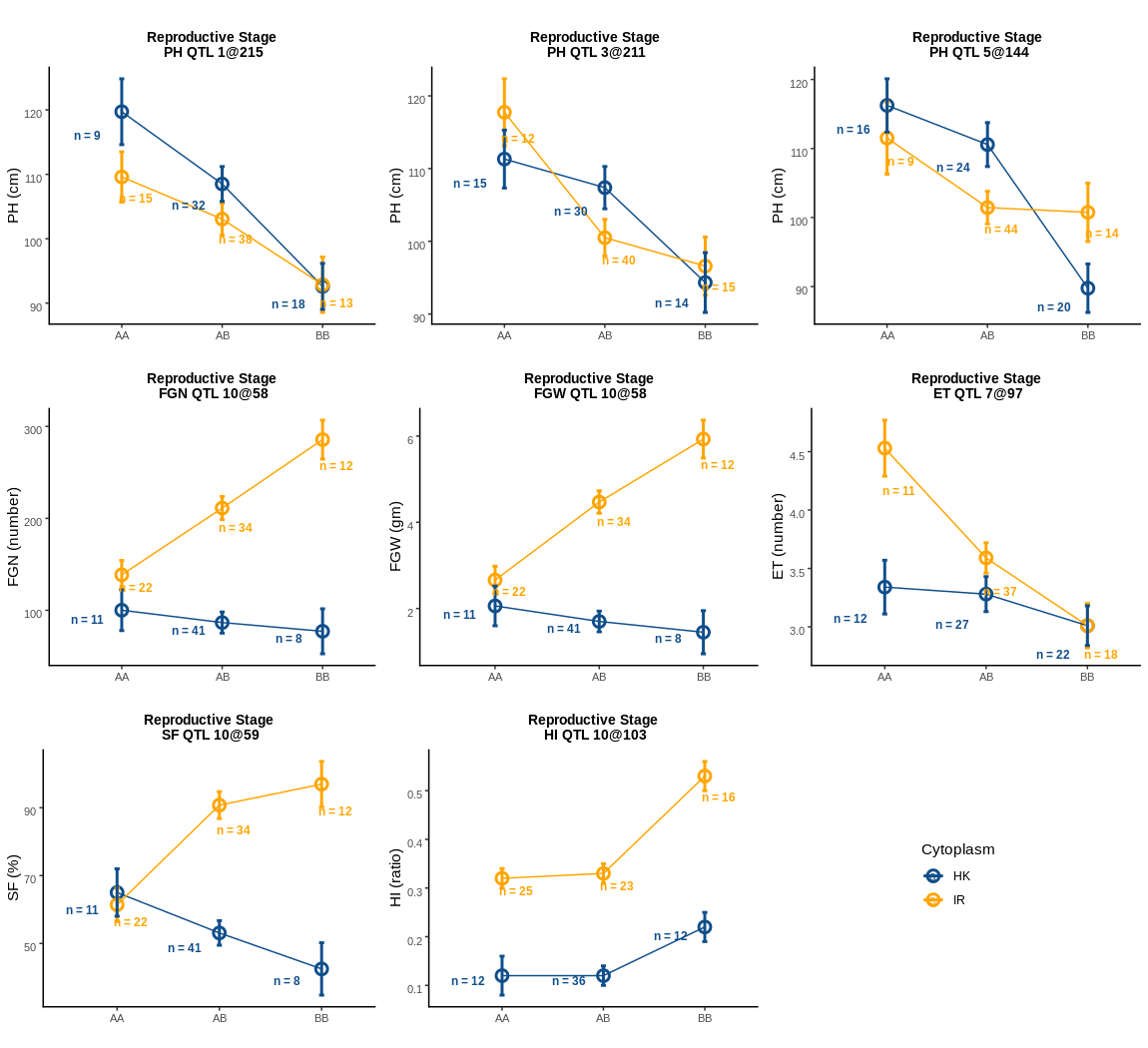

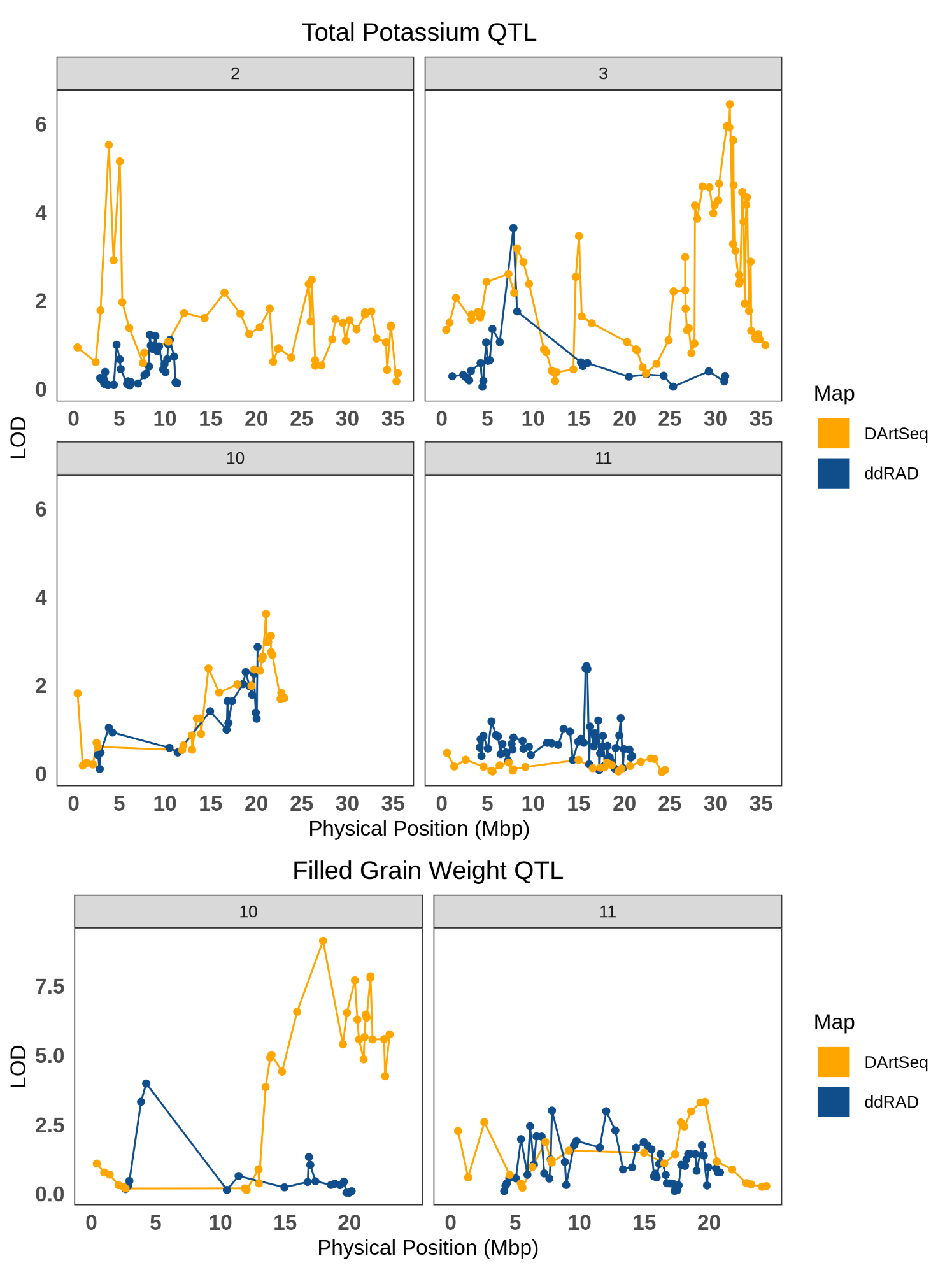

**Noor AUZ, Nurnabi Azad Jewel GM, Haque T, Elias SM, Biswas S, Rahman MS, Seraj ZI**. 2019. Validation of QTLs in Bangladeshi rice landrace Horkuch responsible for salt tolerance in seedling stage and maturation. Acta Physiologiae Plantarum **41**, 173.
